# Immune Response following FLASH and Conventional Radiation in Diffuse Midline Glioma (DMG)

**DOI:** 10.1101/2022.11.09.515507

**Authors:** Oscar Padilla, Hanna E. Minns, Hong-Jian Wei, Weijia Fan, Andrea Webster-Carrion, Masih Tazhibi, Nicholas M. McQuillan, Xu Zhang, Matthew Gallitto, Rebecca Yeh, Zhiguo Zhang, Tom K. Hei, Luca Szalontay, Jovana Pavisic, Yuewen Tan, Naresh Deoli, Guy Garty, James H. Garvin, Peter D. Canoll, Claire Vanpouille-Box, Vilas Menon, Marta Olah, Raul Rabadan, Cheng-Chia Wu, Robyn D. Gartrell

**Affiliations:** Department of Radiation Oncology, Columbia University Irving Medical Center, New York, NY; Department of Pediatrics, Columbia University Irving Medical Center, New York, NY; Mailman School of Public Health, Columbia University, New York, NY; Institute for Cancer Genetics, Columbia University Irving Medical Center, New York, NY; Department of Genetics and Development, Columbia University Irving Medical Center, New York, NY; Center for Radiological Research, Columbia University Irving Medical Center, New York, NY; Radiological Research Accelerator Facility, Columbia University Irving Medical Center, Irvington, NY; Department of Pathology and Cell Biology, Columbia University Irving Medical Center, New York, NY; Department of Radiation Oncology, Weill Cornell Medicine, New York, NY; Department of Neurology, Columbia University Irving Medical Center, New York, NY; Center for Translational and Computational Neuroimmunology, Columbia University Irving Medical Center, New York, NY; Taub Institute for Research on Alzheimer’s Disease and the Aging Brain, Columbia University Irving Medical Center, New York, NY; Department of Systems Biology, Columbia University Irving Medical Center, New York, NY; Department of Biomedical Informatics, Columbia University Irving Medical Center, New York, NY; Program for Mathematical Genomics, Columbia University Irving Medical Center, New York, NY; Department of Radiation Oncology, Icahn School of Medicine at Mount Sinai, New York, NY; Department of Oncology, Division of Pediatric Oncology, Johns Hopkins School of Medicine, Baltimore, MD; Oregon Health and Science University School of Medicine Portland, OR

**Keywords:** FLASH, radiation, diffuse midline glioma, DMG, DIPG, tumor microenvironment, immune modulation, genome biology, single cell technologies

## Abstract

**Purpose:** Diffuse Midline Glioma (DMG) is a fatal tumor traditionally treated with radiotherapy (RT) and previously characterized as having a non-inflammatory tumor immune microenvironment (TIME). FLASH is a novel RT technique using ultra-high dose rate, which is associated with decreased toxicity and effective tumor control. However, the effect of FLASH and conventional (CONV) RT on the DMG TIME have not yet been explored.

**Methods:** Here, we perform single-cell RNA sequencing and flow cytometry on immune cells isolated from an orthotopic syngeneic murine model of brainstem DMG following the use of FLASH (90Gy/sec) or CONV (2Gy/min) dose-rate RT, and compare to unirradiated tumor (SHAM).

**Results:** At day 4 post-RT, FLASH exerts similar effects as CONV in the predominant microglial (MG) population, including the presence of two activated subtypes. However, at day 10 post-RT, we observe a significant increase in type 1 interferon alpha receptor (IFNAR+) in MG in CONV and SHAM compared to FLASH. In the non-resident myeloid clusters of macrophages (MACs) and dendritic cells (DCs), we find increased type 1 interferon (IFN1) pathway enrichment for CONV compared to FLASH and SHAM by scRNA-seq. We observe this trend by flow cytometry at day 4 post-RT in IFNAR+ MACs and DCs, which equalizes by day 10 post-RT. DMG control and murine survival are equivalent between RT dose rates.

**Conclusion:** Our work is the first to map CONV and FLASH immune alterations of the DMG TIME with single-cell resolution. While DMG tumor control and survival are similar between CONV and FLASH, we find that changes in immune compartments differ over time. Importantly, while both RT modalities increase IFN1, we find that the timing of this response is cell-type and dose-rate dependent. These temporal differences, particularly in the context of tumor control, warrant further study.

## INTRODUCTION

Diffuse Intrinsic Pontine Glioma (DIPG), re-classified by the World Health Organization (WHO) in 2021 as Diffuse Midline Glioma, H3K27M-altered (DMG), is a universally fatal pediatric brain tumor that carries a median survival of 9 to 11 months.^1^ ^2^ While radiotherapy (RT) is the standard-of-care treatment for DMG, it only offers a 3 month improvement in survival.^3^ ^4^ Clinical trials have explored a variety of targeted therapies in combination with RT, yet none have prolonged survival.^4^ ^5^.

Combining immunotherapy with RT has improved survival in several adult cancers.^6–9^ However, immunotherapies like immune checkpoint inhibitors have had minimal success in childhood malignancies and combination approaches with RT have not been tried in DMG.^10^ In order to appropriately pursue combination strategies in DMG, we must first understand how RT affects the tumor immune microenvironment (TIME). Previous studies have shown that DMG harbors an immunologically “cold” or non-inflammatory TIME compared to adult glioblastoma (GBM).^11^ ^12^ One study evaluated diagnostic specimens from human DMG and GBM tumors using immunohistochemistry staining and flow cytometry and found that DMGs have less CD3+ T lymphocytes compared to GBM.^11^ This study also used RNA-seq to evaluate macrophages isolated from primary DMG and GBM and found that macrophages from DMG tumors expressed fewer inflammatory factors compared to macrophages from GBM tumors.^11^ Of note, the effect of RT has not been evaluated in human DMG specimens, as biopsies are not performed at this timepoint due to the precarious brainstem location of this tumor. Consequently, a critical gap exists in delineating RT immune responses that may be leveraged in combination with immunotherapy.

FLASH, or ultra-high dose rate RT (>40Gy per second), is a novel technique that demonstrates less tissue toxicity without compromising tumor control when compared to conventional (CONV) dose rate RT (≤2Gy per minute).^13–18^ Specifically, FLASH has been associated with less neurocognitive toxicity compared to CONV, which may be critical when considering RT in pediatric brain tumors.^13^ ^19^ RT at CONV dose rates has been shown to effectively increase the antigenicity of tumor cells via type 1 interferon (IFN1) response.^20^ ^21^ However, CONV can also deplete immune populations and induce regulatory T-cell phenotypes in the TIME.^22^ ^23^. While immune studies are limited with FLASH, previous evidence suggests that when compared to CONV, ultra-high dose rates yield higher CD8^+^ T-cells and myeloid subsets in the TIME of a Lewis lung carcinoma model.^24^ Studies evaluating FLASH in the brain of non-tumor bearing mice find decreases in microglial activation and neuroinflammation when compared to CONV.^24–27^ Moreover, a recent study evaluating proton FLASH in rat glioblastoma found that CONV and FLASH equally increase non-resident immune responses in the TIME when compared to untreated tumor at day 8 post-RT.^27^ Notwithstanding this early evidence, immune responses post-RT in the DMG TIME have not yet been evaluated.

We use single cell RNA sequencing (scRNA-seq) and flow cytometry to analyze the TIME in an immunocompetent syngeneic orthotopic murine model of brainstem DMG post CONV or FLASH and compare to unirradiated tumor (SHAM) and normal brainstem. Our work is the first to map CONV and FLASH immune alterations in the DMG TIME.

## METHODS

### DMG tumor induction

We utilized a syngeneic murine brainstem DMG cell line (PDGFB+, H3.3K27M, p53^-/-^ cell line, 4423 DIPG, Supplemental Methods).^28^ All animal experiments were performed in accordance with national guidelines and approved by our Institutional Animal Care and Use Committee (IACUC). Juvenile 5-week-old immunocompetent male B6(Cg)-Tyrc-2J/J mice were purchased from Jackson Laboratories (Bar Harbor, ME). At 6 weeks of age, mice were anesthetized with isoflurane, immobilized in the stereotactic instrument (Stoelting, Wood Dale, IL, USA), and underwent orthotopic brainstem implantation with 4423 DIPG at 100,000 cells per microliter (uL) per mouse. Tumors were confirmed 10 days post injection (dpi), using MRI (Supplemental Methods).

### Radiation

At 17 dpi, tumor-bearing mice were randomly assigned to receive 15Gy of radiation either at a conventional dose rate (CONV, 2 Gy/min) or at an ultra-high dose rate (FLASH, 90 Gy/sec) using the Ultra-High Dose Rate FLASH irradiator at our experimental irradiator facility, based on a repurposed Varian Clinac 2100C.^29^ Mice were anesthetized using isoflurane and immobilized using a device incorporating 6.4mm thick adjustable lead shielding, which was aligned to external anatomy creating an open field around the mouse hindbrain (Supplemental Figure 1). All irradiations were performed using 9 MeV electrons (R_50_=3.9 cm). Further details of radiation methods including beam parameters can be found in Supplemental Methods.

### Experimental design

Mice were evaluated among four groups: CONV (15Gy at 2Gy/min), FLASH (15Gy at 90Gy/sec), no radiation (SHAM), or normal, untreated healthy mice (Normal Brainstem). Four mice were studied per group for a total of 16 mice. Four days post-RT, corresponding to 21 dpi, mice were euthanized and brainstems dissociated for CD45+ cell isolation to probe the immune populations using hashtag scRNA-seq^30^ (Figure 1). Flow cytometry was also performed separately in mice from CONV, FLASH, and SHAM groups at day 4 (D4) and day 10 (D10) post-RT, following the same irradiation and experimental protocols.

**Figure 1:**
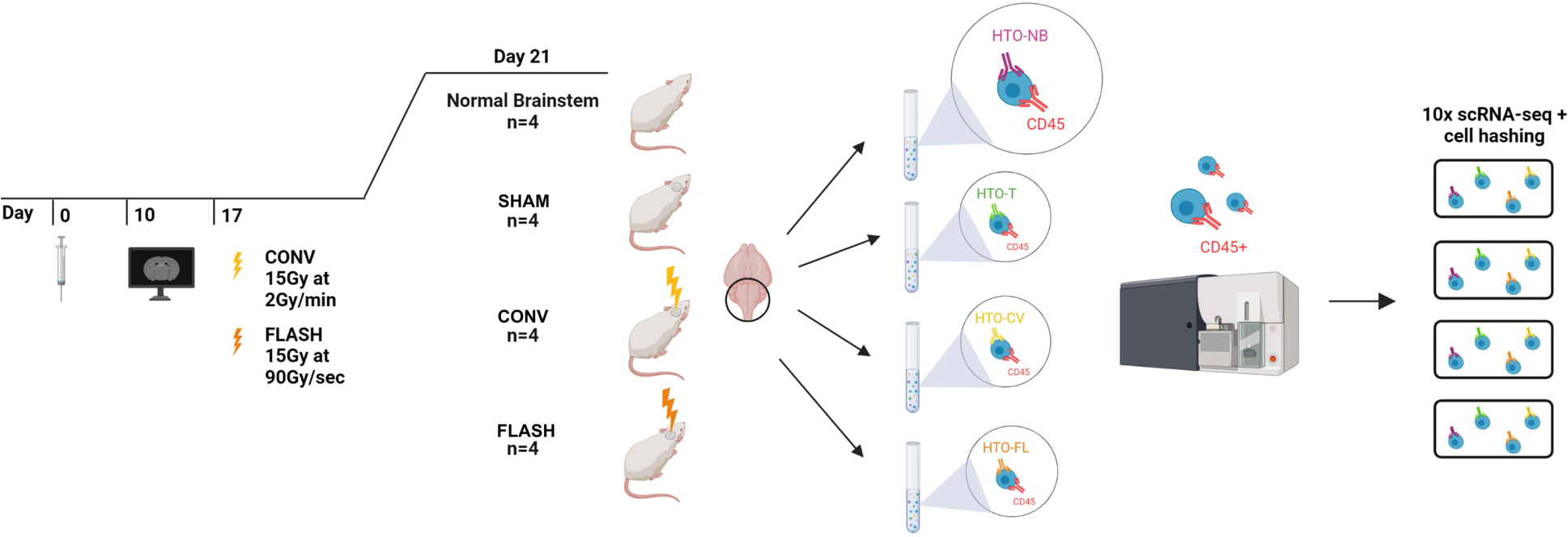
Experimental design for Single Cell Sequencing. 0 days post injection (dpi); H3.3K27M mutant DMG cells are stereotactically injected into the brainstem of juvenile mice. 10 dpi; tumor presence is confirmed by MRI. 17 dpi; mice are randomly assigned to receive either conventional radiation (RT) (CONV, 15Gy at 2Gy/min), FLASH (15Gy at 90Gy/sec), or unirradiated tumor (SHAM). A cohort of 4 untreated, healthy mice are included for analysis of unirradiated brainstems (“Normal Brainstem”). Day 21 (day 4 post-RT); All groups are euthanized for cell isolation, hashtag oligonucleotide (HTO) staining, CD45+ sorting, and finally, single-cell RNA sequencing.

### Immune cell isolation and processing for scRNA-seq

After tissue processing and homogenization (Supplemental Methods), the pellet was resuspended in a staining solution consisting of 7AAD viability solution (Biolegend, 420403), APC anti-mouse CD45 (Biolegend, 147708), and a TotalSeq-B anti-mouse hashtag antibody specific to treatment group (Biolegend; Normal Brainstem:155843, SHAM:155837, CONV:155833, FLASH:155831). The cells were stained for 15 minutes before resuspension in 0.3% BSA (Millipore Sigma, A2058) and transfer to our institution’s Center for Translational Immunology (CCTI) core for CD45 sorting via the Influx cell sorter. The sorting was capped at 30,000 CD45+ cells per mouse and these cells were then brought for sequencing at our institution’s Genome Center’s Single Cell

### Analysis Core

Four samples, one sample from each treatment group, were loaded onto a single channel of a 10x Chromium chip. Viability of each sample was assessed to ensure an equal number of viable cells were taken from each sample then included in the pooled sample containing with a total of 10,000 cells, which were submitted for scRNA-seq (Supplemental Methods).

### Flow Cytometry

Following the same tissue dissection, homogenization and isolation protocol as done for scRNA-seq, mice were euthanized at D4 post-RT (21 dpi) and D10 post-RT (27 dpi). Cells were stained with a total of 14 markers including 7AAD, CD45, CD11B, CD163, CD206, CD11C, MHC-II, CD3, CD8, CD4, GR1, TMEM119, CD19 and IFNAR1 (Supplemental Methods). Antibody combinations against the following targets were used to define cell types (Supplemental Figure 3): microglia (MG), CD45loCD11B+GR1-TMEM119+; macrophages (MACs), CD45hiCD11B+GR1-TMEM119-; myeloid derived suppressor cells (MDSCs), CD45hiCD11B+GR1+TMEM119-; dendritic cells (DCs), CD45+CD11C+MHCII+; CD19+ B-cells, CD45hiCD11B-CD3-CD19+; T-Cells, CD45HiCD11B-CD3+; CD8+ T-Cells, CD45hiCD11B-CD3+CD8+CD4-; CD4+ T-Cells, CD45hiCD11B-CD3+CD8-CD4+. These antibody combinations were chosen based on previous studies.^31–33^

### Statistical Analysis

Statistical analysis was performed using GraphPad Prism 9.1.2 (GraphPad Software, Inc.). Mean tumor volumes at 15 dpi and 24 dpi (7 days post-treatment) were compared in mice treated with CONV (n = 5), FLASH (n = 5) and SHAM (n = 5) using 2way ANOVA for repeated measures with Greenhouse–Geisser correction and multiple comparisons correction using Tukey. Survival analysis was performed using log-rank Mantel–Cox test comparing median survival times between CONV (n = 4), FLASH (n = 5) and SHAM (n = 4). Single cell analysis processing and statistical analysis are detailed in Supplemental Methods. Flow cytometry data was compared between each group at D4 and D10 separately (n = 5 mice per group at each time point) using Kruskal-Wallis test. Post-hoc pairwise comparison was conducted using Dunn’s test without multiple comparison adjustment. Within each group, comparisons of D4 vs D10 was done using Mann Whitney U Test. Significance was defined as p value of ≤0.05.

## RESULTS

### FLASH and CONV confer isoeffective DMG control

To assess the effect of CONV and FLASH on tumor control we evaluated tumor volume between pre- and post-RT timepoints using MRI and observed a significant decrease in tumor volume at day 7 post-RT (24 dpi) after both CONV (p=0.0006) and FLASH (p=0.0008) compared to SHAM (Figure 2). Further, we performed survival analysis with log-rank Mantel–Cox test and find no significant differences in median survival times between groups (CONV median survival = 28 days, FLASH = 30 days, SHAM = 28 days, p=0.2995; Supplemental Figure 2).

**Figure 2:**
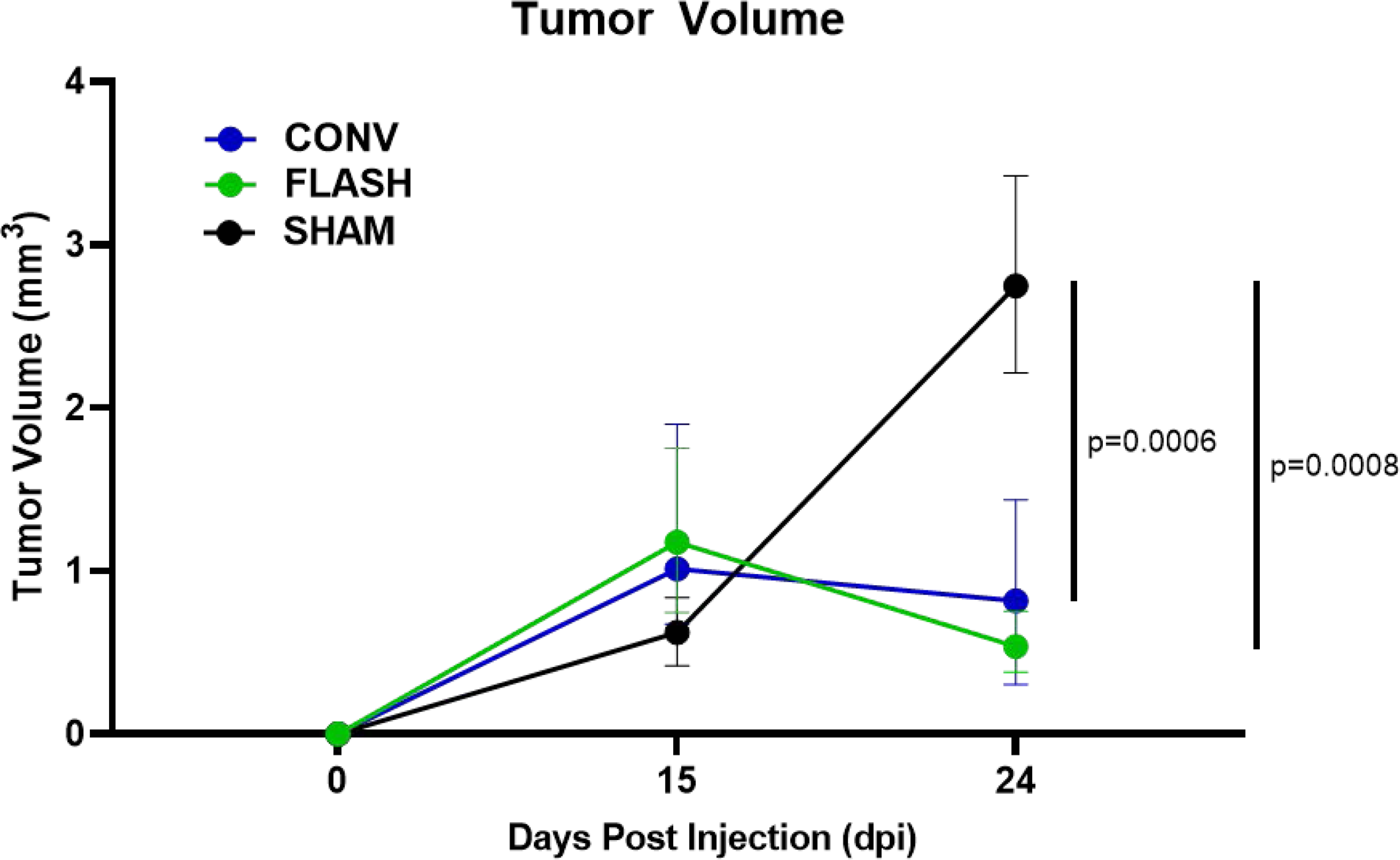
FLASH and CONV tumor volume comparisons. Tumor volume analysis for FLASH (n = 5), CONV (n = 5), and SHAM (n = 5) mice measured at 15 days post injection (dpi) and 24 dpi (7 days post-treatment). Statistical analysis performed with 2way ANOVA for repeated measures with Greenhouse–Geisser correction and multiple comparisons correction using Tukey. Significant p-values considered as p < 0.05.

### scRNA-seq identifies unique immune clusters in DMG TIME

Clustering of 33,308 CD45+ cells reveals 17 unique cell subsets (Figure 3a, Supplementary Data 1, 2). We first analyze all clusters in total, agnostic to treatment. From this, we identify four clusters of MG denoted as MG1, MG2, MG3, and MG4. We find three clusters of non-resident myeloid cells including MACs, MONOs and DCs. Of the non-myeloid clusters, the most abundant is a mature B-cell subtype (BC2). The remaining cells are distributed amongst a T-cell cluster (TC), a neutrophil cluster (NEUT), a pre-B-cell cluster (BC1), and a natural killer cell cluster (NK). We also report a mixed population of proliferative cells in a cluster defined by *Mki67* positivity and another mixed population cluster of *S100b+* cells, likely consisting of astrocytes that bypassed CD45 sorting. Lastly, a group of erythrocytes are present based on hemoglobin gene expression (RBC), as well as a cluster of low-quality cells (lq). For individual treatment groups, we examine the proportion of cells coming from each cluster against their total cell number and find variation between groups (Figure 3b, Supplementary Data 1). We perform key marker gene identification for each cell type using differential gene expression analysis, comparing each cluster against all other clusters. From this, we see that each cell type is defined by unique gene signatures and validated by presence of canonical markers (Figure 3c, Supplementary Data 2).

**Figure 3:**
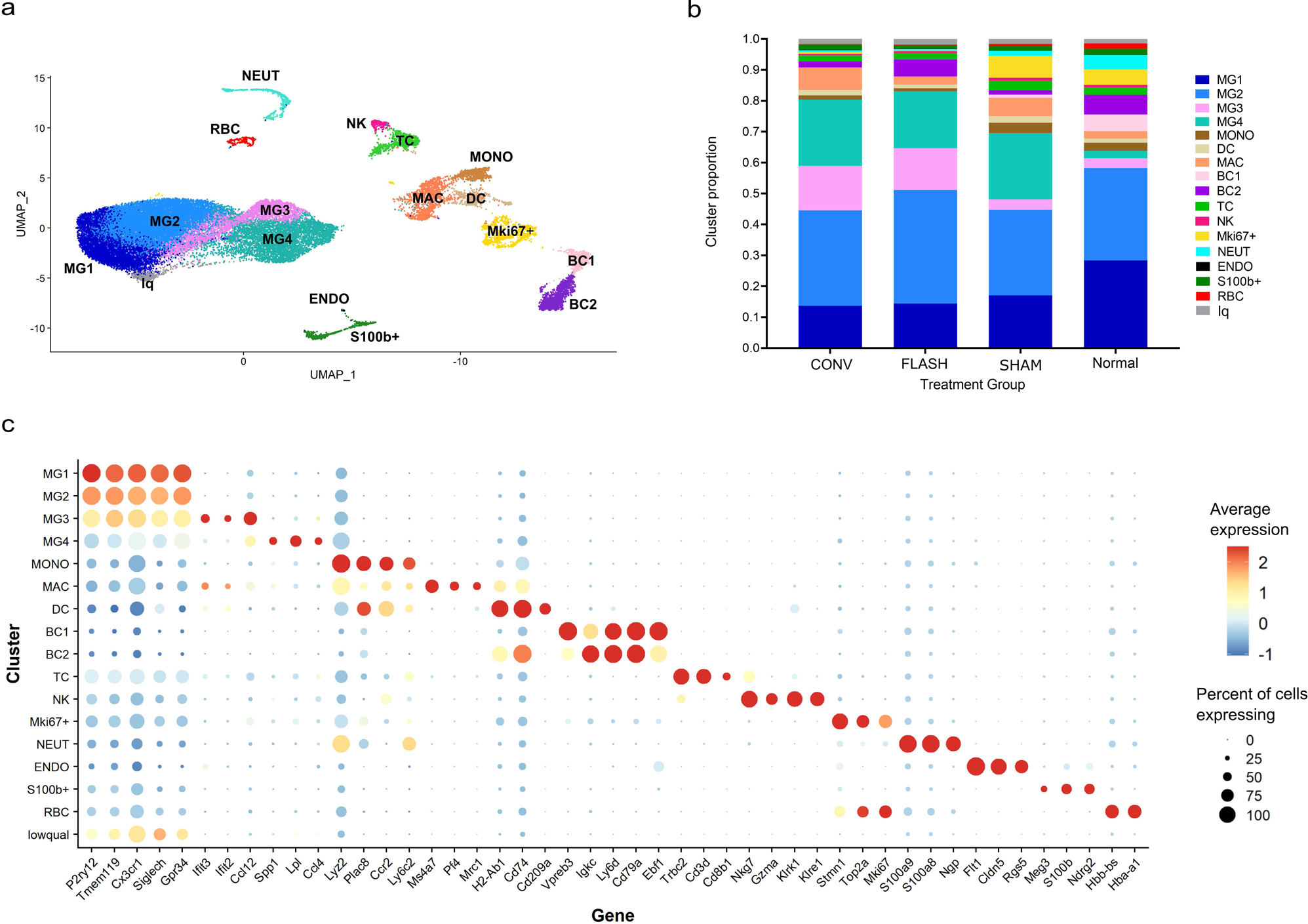
Clustering of CD45+ cells. **(a)** Clustering of 33,308 CD45+ cells reveals 17 unique cell subsets. **(b)** Cluster proportions making up each treatment group when evaluated against the total number of cells collected for that specific group. (c) Key marker genes for each cluster.

### Immune compartment differences between RT and SHAM groups evolve over time

To confirm presence of immune phenotypes and evaluate proportional comparisons between treatment groups, we performed flow cytometry at D4 and D10 post-RT. The gating strategy can be found in Supplemental Figure 3 and all statistical comparisons can be found in Supplementary Data 3. Similar proportions of MG are found at D4 post-RT across CONV, FLASH and SHAM groups (Figure 4a). Further, with the exception of the CONV group in which we observe a significant decrease in MG fraction between D4 and D10 post-RT (p=0.0317), the MG compartment remains insignificantly changed between D4 and D10 post-RT in FLASH and SHAM groups. However, at D10 post-RT there are significantly fewer MG in CONV and FLASH groups compared to SHAM (p=0.0339 and p=0.0109, respectively, Figure 4a). MACs (Figure 4b) are not significantly different at D4 when comparing CONV and FLASH to SHAM. We find a significant fractional increase in MACs from D4 to D10 in the SHAM group (p=0.0079, Figure 4b) while no significant changes are seen in D4 to D10 post-RT in the CONV or FLASH groups. However, at D10, MACs are significantly decreased in CONV (p=0.0196) while in FLASH they trend toward lower proportion compared to SHAM (p=0.0562). With regard to the GR1+ myeloid-derived suppressor cells (MDSCs, Figure 4c), there is no significant difference in proportions at D4 in CONV or FLASH compared to SHAM. However, at D10, CONV and FLASH induce a significant fractional increase relative to SHAM (p=0.0070 and p=0.0086, respectively, Figure 4c), in which MDSCs are observed to decrease between D4 and D10 (p=0.0476, Figure 4c). There are no significant inter- or intra-group differences at D4 or D10 for DCs (Figure 4d). When comparing T-cells (Figure 4e), we see a significant fractional decrease from D4 to D10 in the SHAM group (p=0.0079, Figure 4e), which also demonstrates a significantly lower T-cell fraction relative to CONV and FLASH at D10 (p=0.0072 and p=0.0089 respectively, Figure 4e). We see that these T-cell differences at D10 vary by subtype, with CD8+ T-cells significantly increasing between CONV and SHAM (p=0.0023; Figure 4f) and CD4+ T-cells significantly increasing in FLASH compared to both CONV and SHAM (p=0.0409 and 0.0055, respectively; Figure 4g). Finally, CD19+ B-cells are found to be trending downward from D4 to D10 in CONV (p=0.0556, Figure 4h) and they are significantly increased in SHAM compared to both CONV and FLASH at D10 (p=0.0196 and p=0.0058, respectively, Figure 4h).

**Figure 4:**
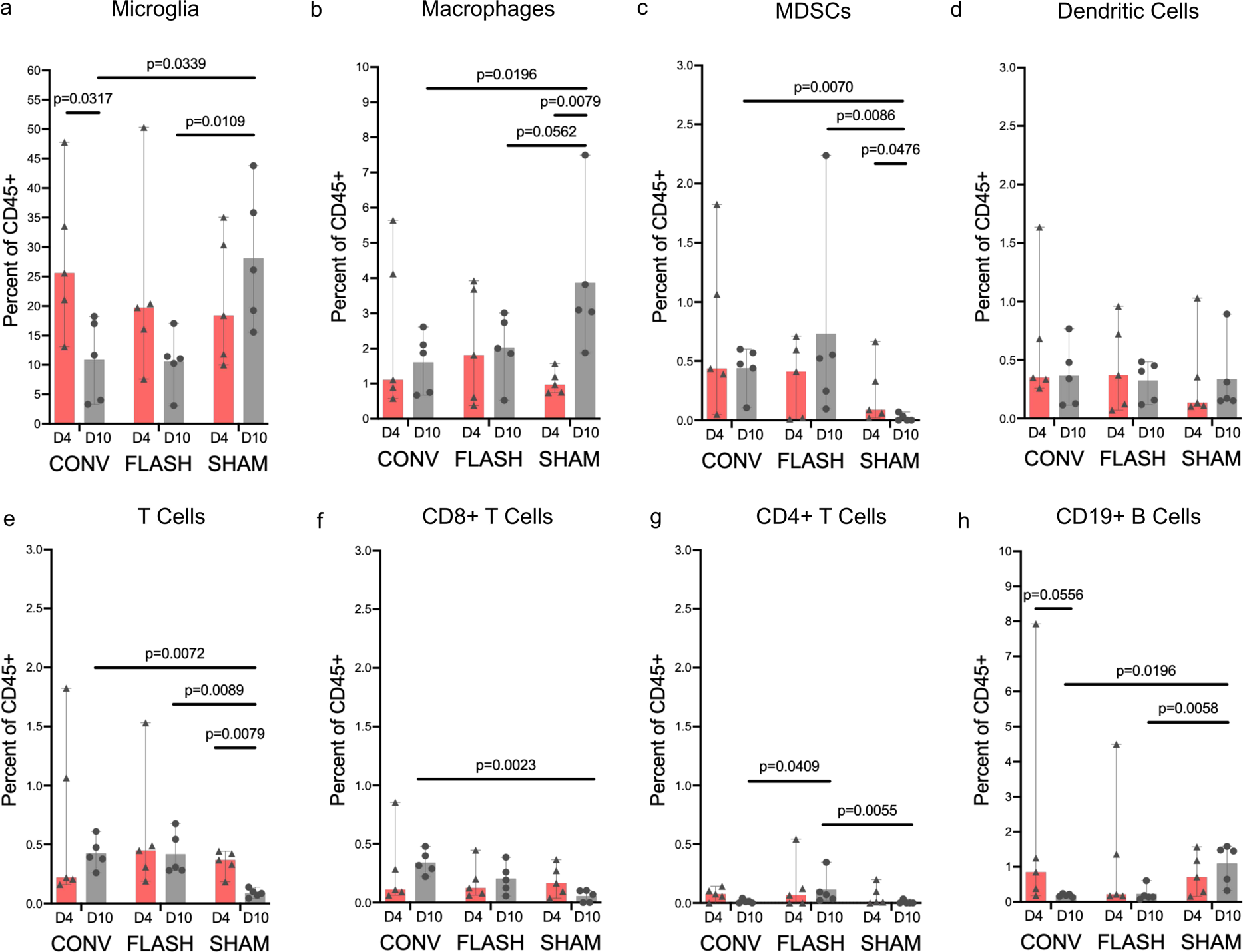
Immune compartment differences as percentage of CD45+ cells comparing CONV, FLASH and SHAM groups (n=5 per time point for each treatment group) at day 4 (D4) and day 10 (D10) post-RT by flow cytometry. (a) microglia, (b) macrophages, (c) myeloid derived suppressor cells (MDSCs), (d) dendritic cells (DCs), (e) CD3+ T Cells, (f) CD8+ T Cells, (g) CD4+ T Cells, (h) CD19+ B Cells. Comparisons of each group at D4 and D10 separately was done using Kruskal-Wallis test with post-hoc pairwise comparison conducted using Dunn’s test without multiple comparison adjustment to provide p-values here. Within each group, comparisons of D4 vs D10 was done using Mann Whitney U Test. Statistical data available in Supplementary Data 3 with significant p-values considered as p ≤ 0.05 highlighted in green and near significant <0.1 yellow.

### MG annotation reveals four distinct subtypes via scRNA-seq

MG represent the predominant population on scRNA-seq. MG are defined by expression of canonical genes such as *P2ry12*, *Siglech* and *Tmem119*,^33^ ^34^ and resolve into four distinct subtypes (MG1, MG2, MG3, and MG4) based on unique gene sets (Figure 5a). Expression of homeostatic marker *P2ry12* decreases from MG1 to MG4, while expression of *Apoe*, a marker of MG activation, increases in the MG3 and MG4 subtypes (Figure 5b).^33^ ^35^ MG1 and MG2 share similar profiles (Supplementary Data 2). MG3 is enriched for multiple interferon genes such as *Ifit3* and *Isg15* (Figure 5c, Supplementary Data 2) while MG4 shows high expression of metabolism-associated genes such as *Lpl* (lipoprotein lipase) and *Fabp5* (fatty-acid binding protein) (Figure 5d, Supplementary Data 2). We ran the top 50 differentially expressed genes in MG3 through REACTOME for pathway analysis (Supplemental Methods), with the top 3 pathways consisting of interferon alpha/beta signaling, interferon signaling, and cytokine signaling (Supplemental Figure 4a). We ran the top 50 genes from MG4 through REACTOME, with the most significant pathways consisting of neutrophil degranulation, interleukin-10 signaling, glucose metabolism, and iron uptake (Supplemental Figure 4b).

**Figure 5:**
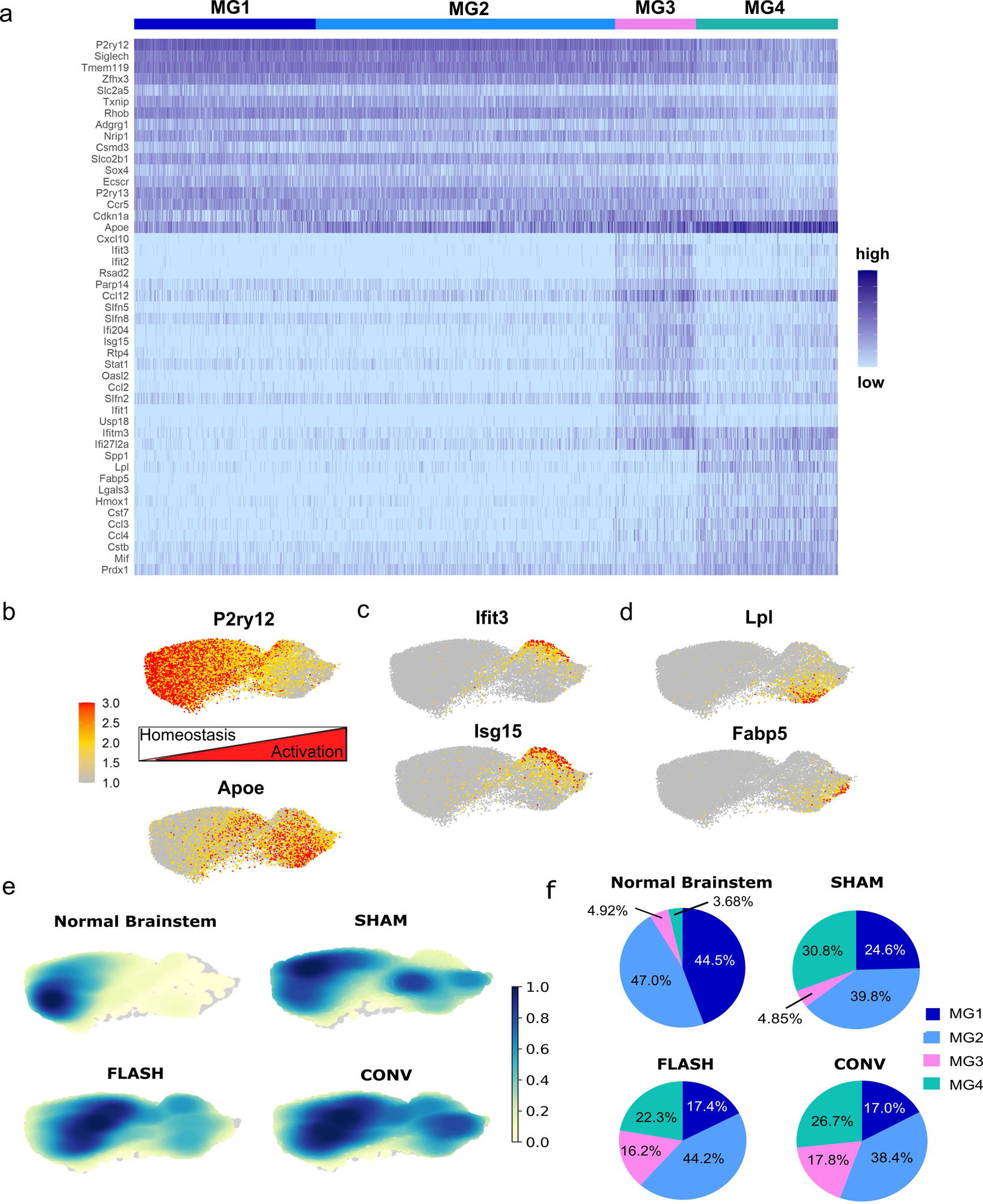
Characterization and localization of MG subtypes across treatment groups. **(a)** Heatmap of top marker genes from each MG cluster. Feature plot of average expression of **(b)** P2ry12 and Apoe **(c)** Ifit3 and Isg15 and **(d)** Lpl and Fabp5 across the MG clusters. **(e)** Density of cells from each treatment group visualized across MG clusters. **(f)** MG cluster proportions within each treatment group, determined as the fraction of MG subtype cells over the total number of MG cells collected for that specific group.

In Normal Brainstem, MG1 and MG2 account for the majority of MG (44.5% and 47.0% respectively), with less than 10% of cells present in MG3 and MG4 clusters (Figure 5e, f). With SHAM, there is an expansion of MG4 (30.8%) while MG3 (4.85%) is maintained at a similar proportion as in Normal Brainstem (Figure 5e, f). Upon irradiation with either CONV or FLASH, the MG3 cluster undergoes significant expansion (17.0% and 16.2% respectively), while MG4 experiences moderate reduction (26.7% and 22.3% respectively) compared to SHAM (Figure 5e, f).

### FLASH and CONV induce similar MG subtypes and early IFN1 response

We next perform two individual analyses to identify genomic alterations uniquely induced by FLASH and by CONV relative to SHAM: FLASH vs SHAM (FvS) and CONV vs SHAM (CvS), respectively (Supplementary Data 4). We compare both the common and the different up- and down-regulated genes resulting from each analysis. For all MG clusters, both RT modalities upregulate DNA damage response genes associated with ionizing radiation (i.e. *Cdkn1a*, *Phlda3*, *Bax*) to a similar extent (Supplemental Figure 4c, e).^36^ ^37^ In MG1 and MG2, FLASH and CONV similarly downregulate innate immunity chemokines *Ccl9* and *Ccl6* compared to SHAM (Supplemental Figure 4c). In MG3, both RT interventions similarly downregulate *Ccl9* along with *Klf2* (Supplemental Figure 4e), a myeloid repressor of neuroinflammation.^38^ In MG4, both RT groups similarly upregulate IFN1 genes such as *Isg15, Ifitm3*, and *Ifit3*, while decreasing expression of major histocompatibility complex (MHC) class II genes *H2-Aa* and *H2-Ab1* compared to SHAM (Supplemental Figure 4e). Regarding differential upregulation of genes, FLASH shows unique enrichment for *Apoe* (MG1, MG3) and *Ccl12* (MG1, MG2), while CONV shows distinct increases in *Cd52, Ifi27l2a, Ccl2,* and *Cst7* in MG4 (Supplementary Data 4).

We then ran the most significant, upregulated common genes between FvS and CvS for each MG cluster through REACTOME and find pathways associated with cell cycle checkpoints and DNA damage response (Supplemental Figure 4d, f). For MG4 specifically, we also find upregulation of common pathways relating to interferon alpha/beta signaling and cytokine signaling (Supplemental Figure 4f). Using flow cytometry, we stained MG with IFNAR to evaluate IFN1 response. Representative IFNAR peak shifts for the relevant cell types can be found in Supplemental Figure 3 and all statistical comparisons can be found in Supplementary Data 3. We find that at D4, there is no significant difference in IFNAR+ MG between FLASH and CONV compared to SHAM (Supplemental Figure 5a). At D10, the percentage of IFNAR+ MG decreases in the FLASH treated mice relative to CONV (p=0.0162) and SHAM (p=0.0403) with no significant differences in intra-group comparisons between D4 to D10 (Supplemental Figure 5a).

### FLASH and CONV induce differential MAC and DC IFN1

We next examine the non-resident myeloid clusters: MONO, MAC, and DC, and find clear segmentation of gene signatures, with varying degrees of overlap in classical myeloid markers such as *Ccr2* and *Vim*^39^ ^40^, and MHC class II genes *H2-Ab1* and *Cd74* (Supplemental Figure 6a). Studying the cell density of these clusters within the non-resident myeloid compartment of each treatment group, we find treatment-specific differences (Supplemental Figure 6b-c). We next perform differential gene expression analysis within the MAC cluster between individual RT groups and SHAM to find common genes associated with both irradiation modalities (FvS and CvS; Supplementary Data 4). We find similarly upregulated genes associated with RT response (e.g. *Cdkn1a, Bax*, and *Phlda3*)^36^ ^37^ as well as similarly downregulated genes associated with antigen presentation (e.g. *Cd74, H2-Eb1, H2-Aa, H2-Ab1*) (Figure 6a). We then perform differential expression analysis to directly compare CONV vs FLASH using a Volcano Plot (Figure 6b, Supplementary Data 4). For CONV, the most upregulated genes compared to FLASH include *Isg15, Irf7, Ifit3, Ifit2, Tgfbi,* and *Tspo* (Figure 6b). For FLASH, the most upregulated genes compared to CONV are *Mrc1, Pf4, Lyve1* and *Cd163* (Figure 6b). Mapping the average expression of these genes onto non-resident myeloid clusters, we find a dichotomy between CONV-(*Isg15 and Ifit3 shown*) and FLASH-associated (*Pf4 and Mrc1 shown*) genes that mirrors their individual cell densities in the MAC cluster (Figure 6c). Considering that the genes enriched in the FLASH group may reflect a particular MAC subtype, we assessed CD163 and CD206 (Mrc1) on MACs by flow cytometry and find no significant differences between FLASH and CONV at D4 or D10 (Supplemental Figure 5b). Importantly, these CD163+CD206+ MACs increase at D10 compared to D4 in both CONV (p=0.0159) and FLASH (p=0.0079) while these cells are not seen at D4 or D10 in SHAM. Moreover, at D10, there are more CD163+CD206+ MACs in both CONV (p=0.0175) and FLASH (p=0.0025) compared to SHAM (Supplemental Figure 5b).

**Figure 6:**
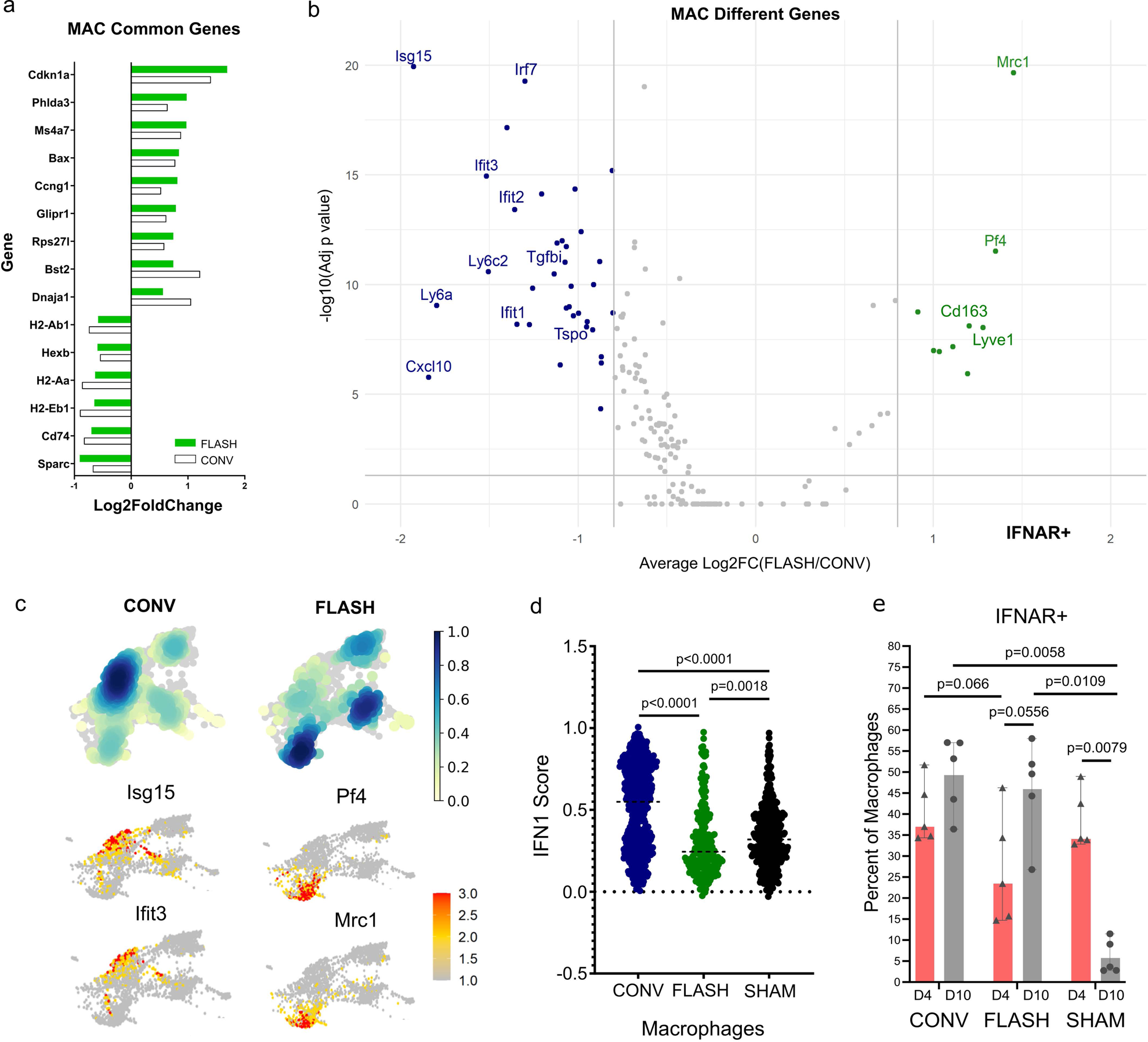
Comparison of immune responses post FLASH and CONV in MACs. **(a)** Common up- and down-regulated genes between independent differential expression analyses of CONV vs SHAM (white) and FLASH vs SHAM (green) in the MAC cluster, plotted by average log2 fold change. **(b)** Volcano plot of the upregulated genes in CONV (left) and FLASH (right) in the MAC cluster. Gray vertical lines indicate average log2 fold change of 0.8 and horizontal gray line indicates adjusted p-value <0.05. **(c)** Density of cells from CONV and FLASH visualized across non-resident myeloid clusters (top). Feature plots of the average expression of top upregulated genes in CONV vs SHAM (left) and FLASH vs SHAM (right) across non-resident myeloid clusters. **(d)** ssGSEA analysis of MACs from CONV, FLASH and SHAM groups using REACTOME interferon alpha/beta gene set (IFN1). **(e)** Percent of IFNAR+ cells per MACs in CONV, FLASH, and SHAM groups at D4 and D10 post-RT. Statistical analysis for flow cytometry data in Supplementary Data 3. Significant p-values considered as p ≤ 0.05.

Next, given the increase in interferon-associated gene expression with CONV, we evaluate IFN1 response by single sample gene set enrichment (ssGSEA) analysis (Supplemental Methods) and find that CONV MACs have a significantly higher IFN1 score compared to FLASH and SHAM (both p < 0.0001; Figure 6d). By ssGSEA, we also find that SHAM MACs have a significantly higher IFN1 score compared to FLASH (p=0.0018, Figure 6d). By flow cytometry, we observe differentially higher IFNAR+ MACs at D4 in CONV vs FLASH that trends toward significance (p=0.066; Figure 4e); this difference disappears at D10, when both RT groups express higher IFNAR+ MAC fractions compared to SHAM (CONV p=0.0058, FLASH p=0.0109). Further, while the fraction of IFNAR+ MACs remains similar between D4 and D10 for CONV, the FLASH fraction trends upward and the SHAM fraction significantly decreases between these timepoints (p=0.0556 and p=0.0079, respectively).

Lastly, we perform differential gene expression analysis in the DC cluster and find only a few genes that are differentially expressed in FvS (upregulated: *Clec4b1, Pid1;* downregulated: *Mdh2, Il1b, Cdk2ap2)* (Figure 7a, Supplementary Data 4). Conversely, CvS has many differentially expressed genes (Figure 7a), with a robust increase in IFN1 related genes such as *Isg15, Irf7, Ifit3,* and *Ifit2,* and significant enrichment of IFN1 score on ssGSEA compared to FLASH and SHAM (both p<0.0001; Figure 7b). We then perform flow cytometry (Figure 7c), and validate our ssGSEA findings, confirming higher IFNAR+ DCs at D4 in CONV vs FLASH and SHAM (both p=0.0196). The percent of IFNAR+ DCs decreases at D10 in CONV (p=0.0079) while FLASH and SHAM remain similar, resulting in no significant differences between treatment groups at D10 (Figure 7c).

**Figure 7:**
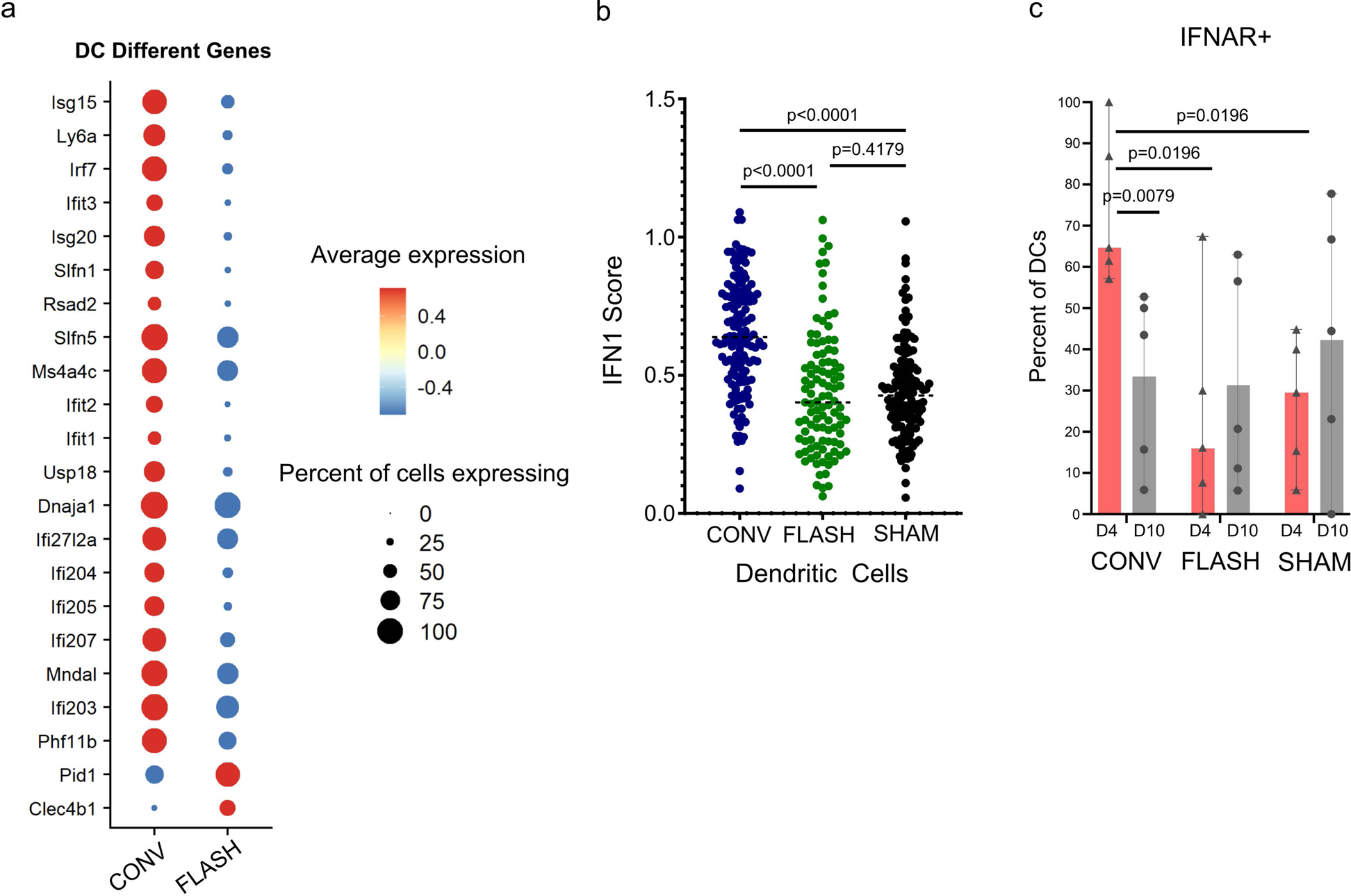
Comparison of immune responses post FLASH and CONV in DCs. **(a)** Dot plot of top 20 most upregulated genes in CONV vs SHAM and the two significantly upregulated genes in FLASH vs SHAM in DCs. **(b)** ssGSEA analysis of IFN1 in DCs from CONV, FLASH and SHAM groups. **(c)** Percent of IFNAR+ cells per DCs in CONV, FLASH, and SHAM groups at D4 and D10. Statistical analysis for flow cytometry data in Supplementary Data 3. Significant p-values considered as p ≤ 0.05.

### Characterization of lymphocyte clusters

NK, T- and B-cell clusters are defined by canonical genes (Supplemental Figure 7a). On FvS and CvS analyses, we find that FLASH upregulates *Lck*, *Klf2, Sell, and Lef1* in T-cells relative to SHAM (Supplemental Figure 7b). These genes are not increased for CONV compared to SHAM. Evaluation by flow cytometry demonstrates higher IFNAR+ T-cells (subtype agnostic) in FLASH vs CONV at D4 only (p=0.0381; Supplemental Figure 7c). Assessing CD8+ T-cells, we see higher IFNAR+ by flow cytometry in CONV vs FLASH at D4 only (p=0.0311; Supplemental Figure 7d). Analysis of B-cells demonstrates BC2 cells are enriched for maturation marker *Ms4a1* and immunoglobulin genes (e.g., *Ighd, Iglc2*)^41^ (Supplementary Data 2). There were few genes differentially enriched for FvS and none for CvS among BC2 cells (Supplementary Data 4). NK cell and BC1 proportions were less than 1% of cells across tumor groups and thus not suitable for differential expression analysis (Supplementary Data 1).

## DISCUSSION

Decades of clinical trials have failed to improve outcomes in DMG, and RT remains the only first-line treatment for this lethal disease. Though RT is known to trigger an anti-tumor immune response, this has not been studied in DMG due to the precarious location of this disease and paucity of available tissue specimens post irradiation.^20^ ^21^ ^42^ Our study aimed to understand the RT immune response of both CONV and FLASH, a novel radiation modality, in the DMG TIME, using a preclinical model that mimics the pathology of human DMG.^28^ Reportedly, FLASH can widen the therapeutic window in high-grade glioma models while also limiting neurocognitive effects and altering the TIME.^15^ ^19^ In this study, we perform high-resolution profiling of the DMG TIME, offering unprecedented insight into the immune imprint of CONV and FLASH RT in this inoperable brainstem tumor.

Consistent with prior FLASH studies in other tumor models, including gliomas, we find equivalent tumor control between dose rates at day 7 post-RT. However, all mice succumbed by day 37 following tumor injection with no survival differences between experimental arms.^15–18^ This is likely a result of the aggressive nature of DMG, which may necessitate a higher biologically effective dose (BED) for durable tumor control than we were able to deliver using a single-fraction regimen.^3^ ^4^ A similar study in a glioblastoma model only achieved survival gains in their RT arms, while maintaining superior cognitive sparing with FLASH, after implementing a hypofractionated schedule with a higher BED than their single-fraction regimen.^15^ Due to logistical reasons, we were only able to administer single-fraction RT at our off-campus animal irradiation facility. Future testing of hypofractionated schedules in DMG is warranted and may further widen the therapeutic window observed with FLASH.

Single-cell technologies have expanded our understanding of the TIME in high-grade gliomas but are limited to pretreatment timepoints.^43^ In our study, we employed scRNA-seq post-RT and resolved 17 unique immune subsets in the DMG TIME. In our model, MG represent the dominant TIME population, and is refined into four subsets reflecting a continuum from homeostatic (MG1 and MG2) to activated phenotypes (MG3 and MG4) with comparable gene expression profiles regardless of RT dose rate. We observe that irradiation with either FLASH or CONV evokes IFN1 in MG, which is consistent with the acute impact of RT in other tumor models and has been previously linked to the recruitment of effector T-cells.^44^ ^45^ We also find that DMG induces a disease-associated MG state in MG4, characterized by hallmark *Apoe, Spp1, Lpl,* and *Fabp5* gene expression and enrichment for interleukin-10 signaling, glucose metabolism and iron uptake pathways.^35^ ^46^ Interestingly, both FLASH and CONV are able to elicit IFN1 expression in MG4, indicating that disease-associated MG retain capacity for immune activation in the setting of RT.

While MG proportions remain grossly similar between dose rates at day 4 by scRNA-seq and flow cytometry, data at day 10 post-RT demonstrates a differential decrease in IFN1 receptor (IFNAR) expression in MG in FLASH compared to CONV and SHAM groups. Prior work shows that CONV can induce an IFN1 response in MG that persists for at least one week post-RT.^37^ In addition, two recent studies demonstrate sustained MG activation at 4 and 10-weeks following CONV, but not FLASH.^25^ ^47^ These findings indicate that FLASH activation of MG is early and transient, and may invoke a different inflammatory cascade than CONV. Whether and how these temporal IFN1 differences in MG are consequential to tumor control and neurocognitive impairment, as has been noted in prior work, remains to be elucidated.^48^ With regard to higher IFNAR^+^ MG in SHAM over FLASH groups at the later timepoint, further investigation is needed to identify the source of this latent IFN1 signaling in the unirradiated TIME. Previous work indicates that cancer cells themselves can produce IFN1 and that this correlates with protumoral signaling in some models.^49^

Compared to MG, we find that non-resident myeloid cells, including MACs, DCs and MONOs, compose less of the DMG TIME. While there is a clear early IFN1 response in infiltrating MACs induced by CONV, this does not manifest in the FLASH group until day 10 post-RT when both FLASH and CONV are higher than SHAM. This finding is opposite to the IFN1 trend observed in MG following FLASH, and may arise from differences in the time kinetics of direct (i.e., resident myeloid activation) versus indirect (i.e., non-resident myeloid recruitment) effects of ultra-high dose rate RT. Moreover, it is possible that the dominance of certain chemotactic cues in the TIME, *Ccl2* in the case of CONV and *Ccl12* for FLASH as shown in our data, leads to the differential recruitment of non-resident myeloid cells. Prior work in traumatic brain injury models demonstrates that *Ccl2* is a more potent chemokine and inflammatory inducer compared to *Ccl12* at acute timepoints.^50^

Overall, we do not see global differences in T-cells between FLASH, CONV and SHAM groups at day 4 post-RT. This could be a result of the early interrogation timepoint, as it may not capture peak T-cell infiltration, which can occur over a broad period of 4 to 10 days following an inflammatory stimulus.^51^ Nevertheless, by day 10 post-RT, we see that CD8^+^ and CD4^+^ T-cells are higher in CONV and FLASH respectively relative to SHAM. Moreover, at this timepoint, CD4+ T-cells are also higher in FLASH relative to CONV. Interestingly, by scRNA-seq, we find that FLASH significantly upregulates genes involved in T-cell activation (*Lck*) and trafficking (*Klf2*), which occurs on the background of higher central-memory (*Sell, Lef1, Klf2*) and naïve (*Sell, Lef1, Ccr7*) T-cell states.^52^ ^53^ However, these markers were not probed by flow cytometry and should be validated on the protein level prior to deriving any conclusions.

Anti-tumor T cell responses can be mediated by DCs via induction of IFN1 post-RT.^20^ ^21^ ^54^ We observe that CONV induces an early IFN1 response in DCs, which is higher than FLASH and SHAM. Although, this does not equate to a differential T-cell infiltrate at day 4 post-RT, we do see that CONV leads to higher IFNAR+ CD8+ T-cells at this timepoint. By day 10 post-RT, we also see that CONV induces a higher CD8+ T-cell infiltrate compared to SHAM; however, this occurs on the background of equivalent IFNAR expression in DCs between these groups.

Understanding the relationship of DCs and T cells in the irradiated TIME, especially along the trajectory of tumor growth or control, is an important next step to improve anti-tumor immune response in DMG. While previous preclinical studies show that FLASH yields higher CD8^+^ T-cells in subcutaneous Lewis lung carcinoma at 6hr post-RT and higher CD4^+^ and CD8^+^ T-cells in orthotopic ovarian cancer at 96hr post-RT, we are limited in drawing direct comparisons with these studies considering the different tumor types and timepoints and the use of a subcutaneous rather than orthotopic model in the first study.^16^ ^24^

Our study has several limitations, including interrogation of the TIME at only two timepoints and flow cytometry analyses focused on some but not all immune subsets identified by scRNAseq (e.g., B-cells subsets). Additionally, we did not include irradiated non-tumor bearing brainstems, which would help to distinguish the immune response arising from irradiation of healthy versus tumor tissue. Future studies should consider inclusion of such control as the diffuse nature of DMG, as observed in our model, complicates the delimitation and separation of healthy versus tumor tissues within a given specimen. The use of a single fraction of 15Gy is also a limitation in our study. While this dose induces an IFN1 response in FLASH and CONV, this does not translate to a survival advantage post-RT, compared to SHAM. As above, this is likely due to the strong resistance pattern of DMG. We selected 15Gy as it is higher than the dose needed to arrest cell proliferation of PDGF-driven DMG *in vitro*, and similar to the normal tissue BED of 14Gy used in a prior FLASH study in orthotopic glioma.^15^ ^55^ Additional FLASH studies in orthotopic glioma models have similarly employed 1 fraction regimens.^15^ ^27^ Future studies focused on hypofractionated dose schedules with more clinically relevant BEDs is an important next step to better understand the effect of RT on the DMG TIME.

In this study, we find that proportions of immune subsets are overall similar between CONV and FLASH, and that both resident and non-resident immune cells in the DMG TIME are able to mount a post-RT IFN1 response. However, there are distinct temporal patterns of IFN1 response within each immune compartment that appear to be dose rate-dependent. The source of these temporal immune alterations, and how these impact tumor control, warrants future investigation.

## Funding

This project was funded by the Hyundai Hope on Wheels Hope Scholar Award (PI: R.D.G, Grant #716838) and Swim Across America, including salary support for R.D.G and H.E.M. O.P. received support from the NCI Stimulating Access to Research in Residency (StARR) Award, supplement to the Columbia Cancer Research Program for Resident Investigators (R38CA231577). C-C.W received support from Gary and Yael Fegel Family Foundation (CU21-1080), St. Baldrick’s Foundation (SBF CU21-0529), the Star and Storm Foundation, Sebastian Strong Foundation, and the Matheson Foundation (UR010590). Development of the FLASH irradiator was partially supported by NIAID Grant U19-AI067773 and by the Departments of Radiation Oncology at Columbia University Irving Medical Center and Weill Cornell Medical Center, as well as by an unrestricted research gift from Barry Neustein. Studies in ZZ laboratory is supported by NINDS (NS 132344). This study used Shared Resources of the Herbert Irving Comprehensive Cancer Center (HICCC) funded in part through Center Grant P30CA013696, specifically Molecular Pathology, Flow Cytometry, Oncology Precision Therapeutics and Imaging (OPTIC), and Genomics and High Throughput Screening Shared Resources. A special thank you to the Flow Cytometry core for their assistance. The content is solely the responsibility of the authors and does not necessarily represent the official views of the NIH.

## Acknowledgements

We would like to thank Dr. Oren J. Becher for providing the murine DIPG 4423 cell line, and Jim Sharkey, Ron Drake and James Viera for their assistance with the FLASH irradiator including Clinac setup and maintenance. Additionally, we would like to thank the Radiological Research Accelerator Facility (RARAF) facility at Columbia University for their help with the FLASH irradiation experiments.

## Conflict of Interest Statement

We have no declarations to report.

## Data Sharing Statement

Demultiplexed sequencing data have been deposited in Synapse under project name, “FLASHvCONV_DMG” (Project synID: syn53166057). Raw data available upon request to corresponding author.

## Statistics

Dr. Robyn Gartrell is responsible for all statistical analyses with support from Weijia Fan.

## Notes

### Competing Interest Statement

The authors have declared no competing interest.

### Summary of Updates

Adding statistic methods and altered discussion. Adding authors.

