## Supplementary figures and images for "Immune Response following FLASH and Conventional Radiation in Diffuse Midline Glioma (DMG)"

### Supplemental Figure 1

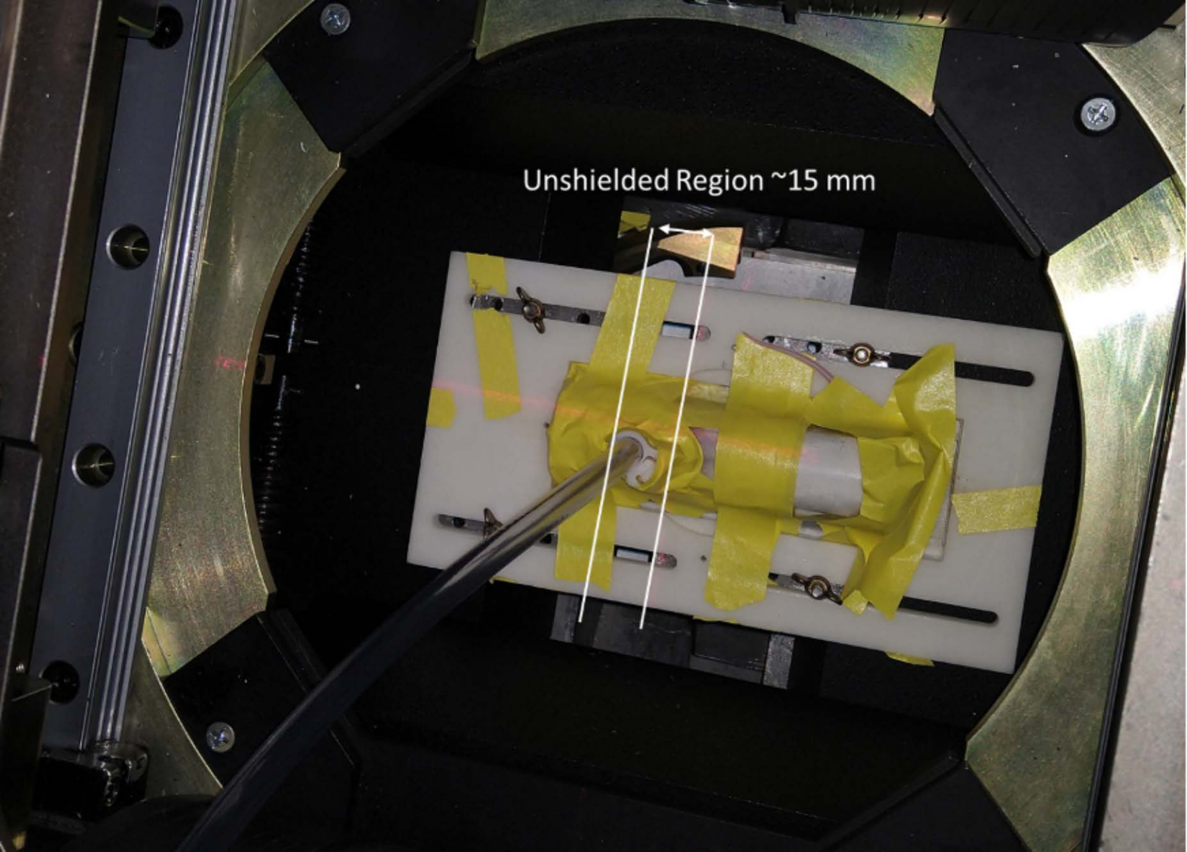

### Supplemental Figure 2

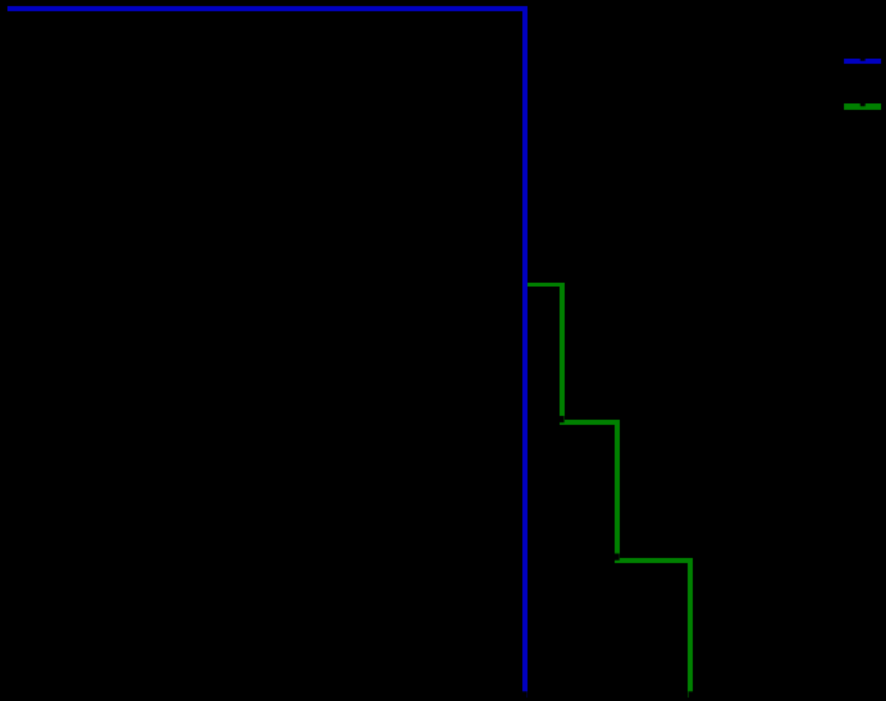

### Supplemental Figure 3

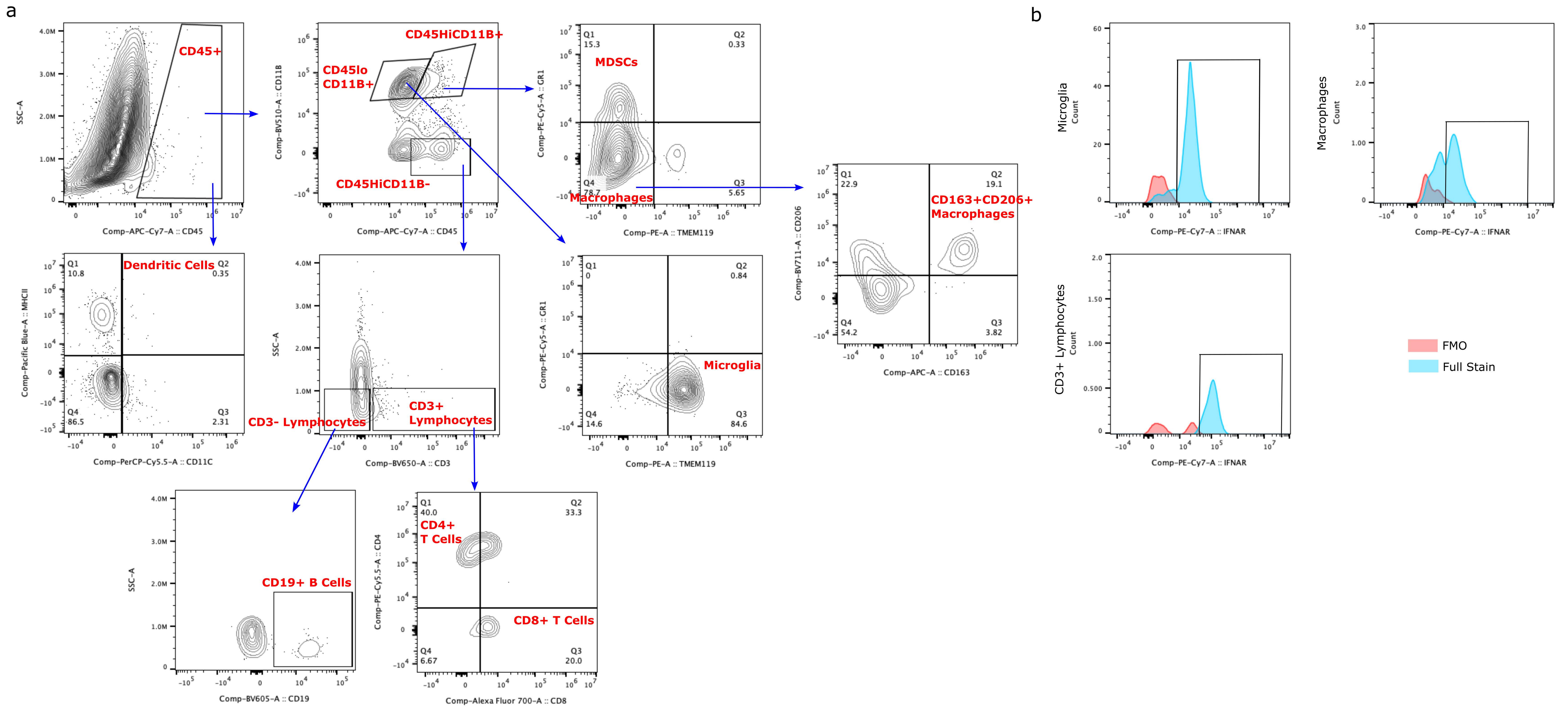

### Supplemental Figure 4

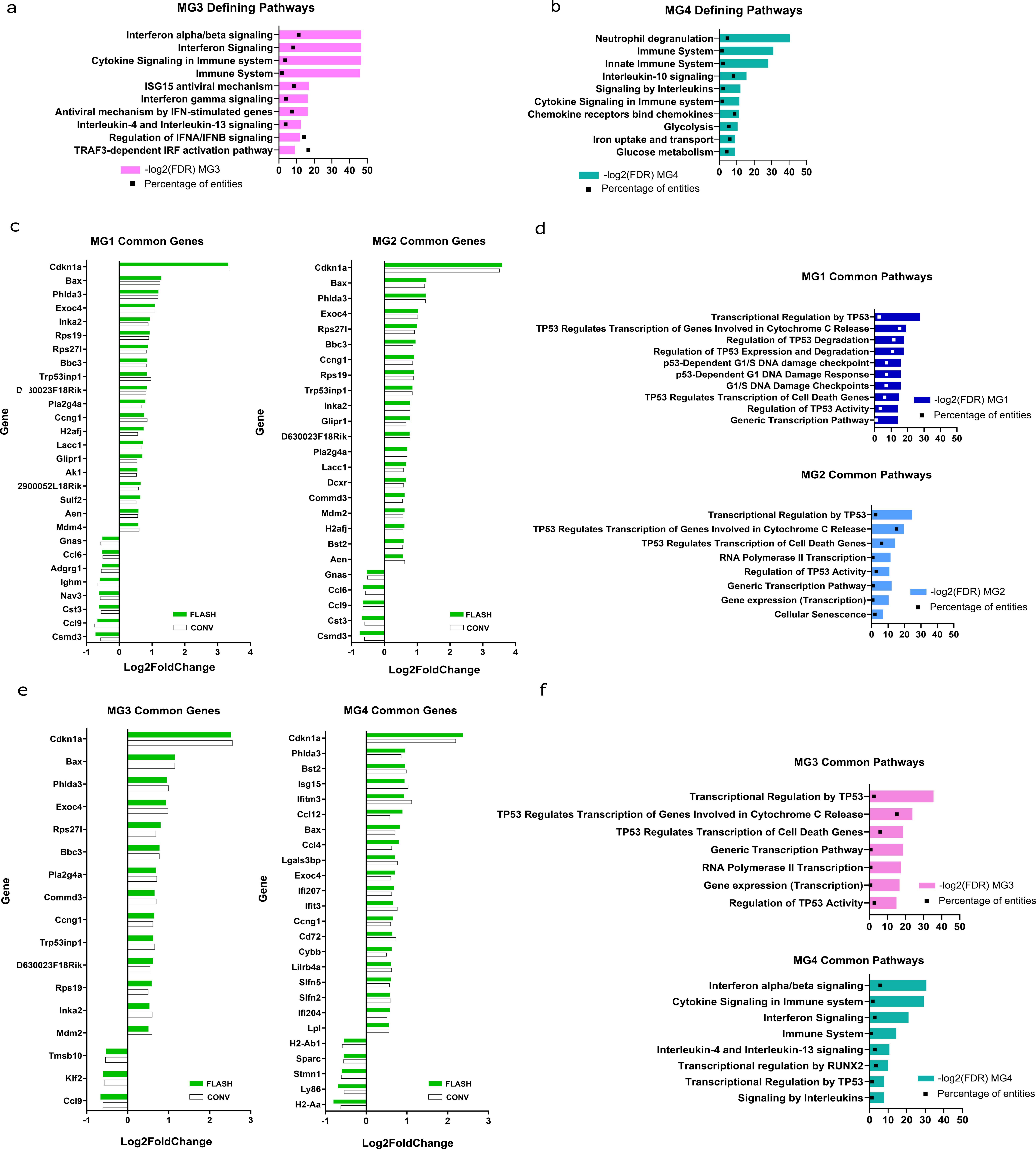

### Supplemental Figure 5

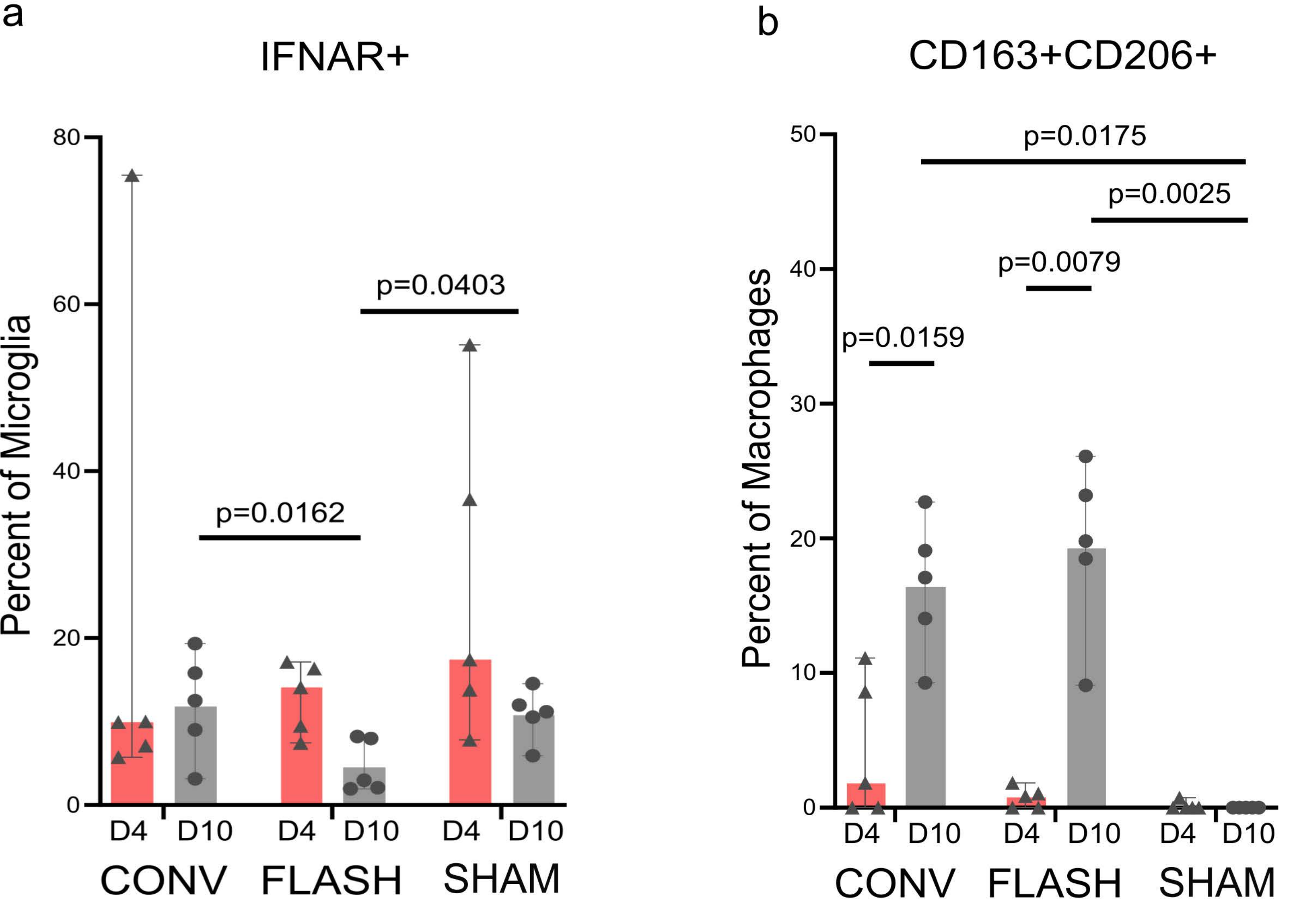

### Supplemental Figure 6

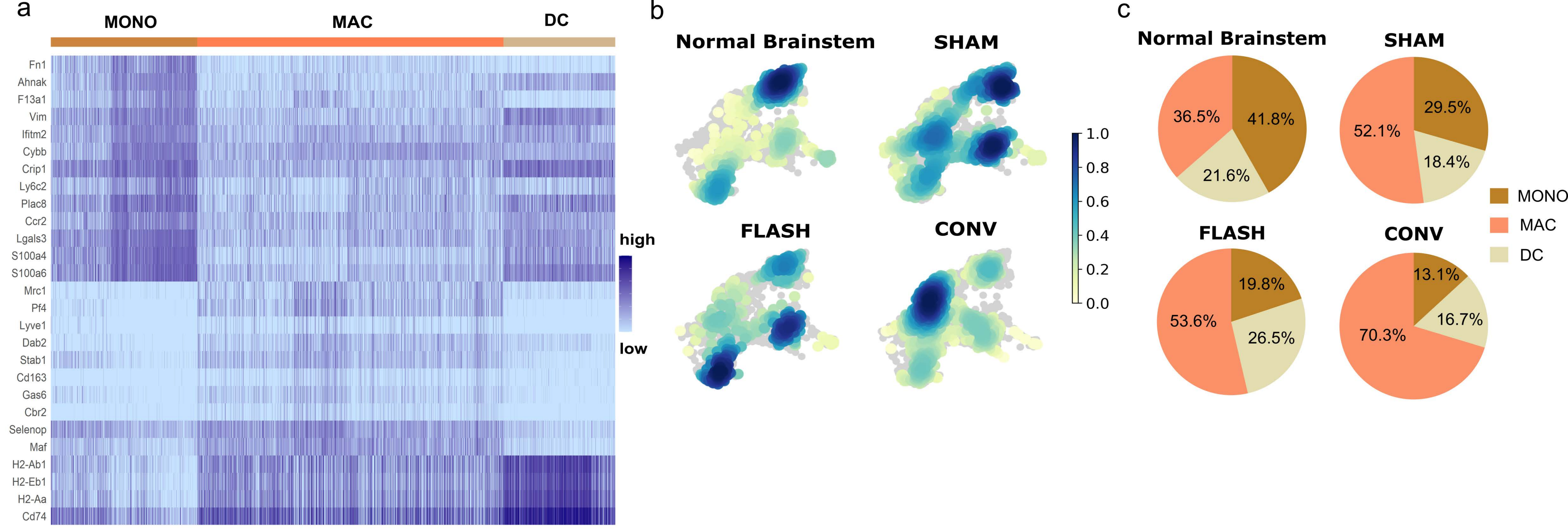

### Supplemental Figure 7

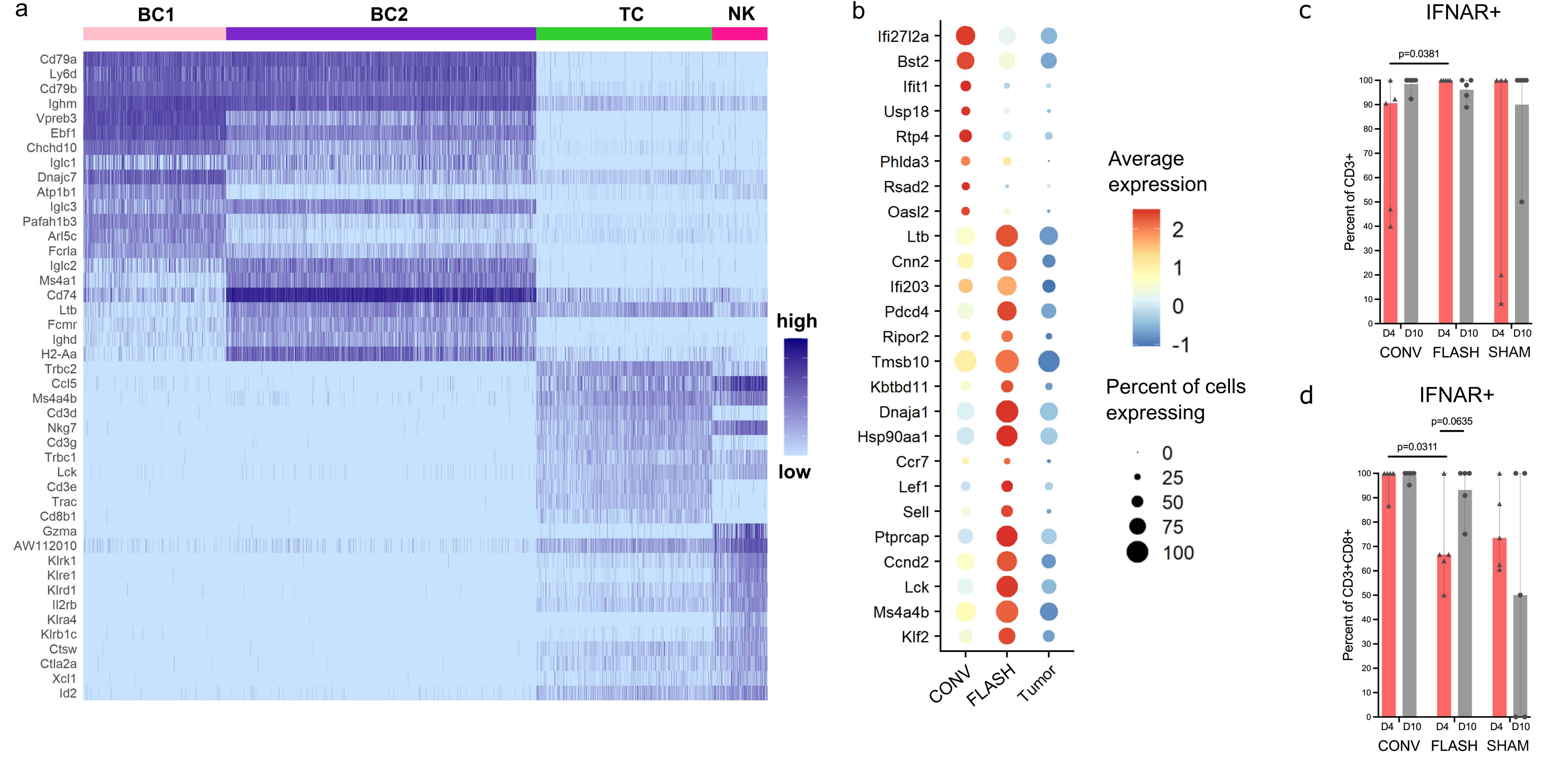
