## Supplemental Figure Legend for "Immune Response following FLASH and Conventional Radiation in Diffuse Midline Glioma (DMG)"

### SUPPLEMENTAL FIGURE LEGENDS

**Supplemental Figure 1: RT jig with unshielded region corresponding to mouse hindbrain area.**

**Supplemental Figure 2**: **Survival Comparisons.** Survival analysis of mice treated with CONV (n = 4, median = 28 days), SHAM (n = 4, median = 28 days) and FLASH (n = 5, median = 30 days). Log rank p=0.2995.

**Supplemental Figure 3: Flow cytometry gating strategy. (a)** Representative gating strategy for major cell types discussed (labelled in red) and CD163+CD206+ macrophage subtype. **(b)** Representative IFNAR1 peak shifts and gating for discussed cell types comparing fluorescence minus one (FMO, lacking IFNAR1 antibody) sample to all stain sample.

**Supplemental Figure 4:** Further annotation and comparisons of microglia subtypes. REACTOME pathway analysis using the top 50 genes showing the 10 most significant pathways (a) from MG3 (b) from MG4. (c) Common up- and down-regulated genes between the independent differential expression analyses of CONV vs SHAM (white) and FLASH vs SHAM (green) in MG1 and MG2, plotted by average log2 fold change. (d) REACTOME pathway analysis of the common upregulated genes between the independent differential expression analyses of CONV vs SHAM and FLASH vs SHAM in MG1 (dark blue) and MG2 (light blue). (e) Common up- and down-regulated genes between independent differential expression analyses of CONV vs SHAM (white) and FLASH vs SHAM (green) in MG3 and MG4, plotted by average log2 fold change. (f) REACTOME pathway analysis of common upregulated genes between independent differential expression analyses of CONV vs SHAM and FLASH vs SHAM in MG3 (pink) and MG4 (turquoise). For REACTOME pathway analysis (a,b,d,f), significance of the pathway analysis is reported using–log2 of the false discovery rate (FDR) and the percentage of entities found in the input gene list compared to the total number of genes in each pathway.

**Supplemental Figure 5: Additional flow cytometry subtype comparisons. (a)** Percent of IFNAR+ cells per microglia in CONV, FLASH, and SHAM groups at D4 and D10. **(b)** Percent of CD163+CD206+ cells per macrophages in CONV, FLASH, and SHAM groups at D4 and D10. Statistical analysis for flow cytometry data in Supplemental Data 3. Significant p-values considered as $p < 0.05$.

**Supplemental Figure 6: Characterization and localization of treatment groups across non-resident myeloid clusters. (a)** Heatmap of top marker genes representing monocytes (MONO), macrophages (MAC), and dendritic cells (DC). **(b)** Density of cells coming from each treatment group visualized across non-resident myeloid clusters. **(c)** Non-resident myeloid subtype proportions within each treatment group, determined as the fraction of subtype cells over the total number of non-resident myeloid cells collected for that specific group.

**Supplemental Figure 7: Annotation of lymphocyte clusters and evaluation of FLASH and CONV immune responses including IFN1 within T Cells. (a)** Heatmap of the top marker genes from each lymphocyte cluster. **(b)** Dot plot of the top upregulated genes in CONV-RT vs SHAM and FLASH vs SHAM in T-cells. **(c)** Percent of IFNAR+ cells per CD3+ T Cells in

CONV, FLASH, and SHAM groups at D4 and D10. **(d)** Percent of IFNAR+ cells per CD3+CD8+ T Cells in CONV, FLASH, and SHAM groups at D4 and D10. Statistical analysis for flow cytometry data in Supplemental Data 3. Significant p-values considered as $p < 0.05$.

**Supplemental Figure 8: Clustering based on our four individual runs on the 10X genomics chromium, grouped both by cell type and treatment group, before combining the datasets.**

**Supplementary Data 1: scRNAseq cell proportions per group and per cluster**

**Supplementary Data 2: Key marker gene list for each cluster.**

**Supplementary Data 3: Flow cytometry data and statistics.**

**Supplementary Data 4: Differential expression analysis comparing FLASH vs SHAM and CONV vs SHAM for each cluster.**
