## Supplemental Methods for "Immune Response following FLASH and Conventional Radiation in Diffuse Midline Glioma (DMG)"

### SUPPLEMENTAL METHODS WITH REFERENCES

#### DIPG tumor cell culture

We utilized a previously published syngeneic murine brainstem DMG cell line (PDGFB+, H3.3K27M, $p53^{-/-}$ cell line, 4423 DIPG).[1] Cells were grown at 37 degrees and 5% $CO_2$ in mouse neurocult media (Stem cell Technologies, 05700) containing 10% mouse cell proliferation supplement (Stem Cell Technologies, 05701), 1% Pen-Strep (Invitrogen, 15140-122), 20ng/ml Human basic FGF (Invitrogen, 13256-029), 10ng/ml Human EGF (Invitrogen, PHG0314), and 2ug/ml Heparin (Stem Cell Technologies, 07980).

#### DMG tumor cell preparation for tumor induction

4423 DIPG Cells were prepared in suspension with DMEM (Dulbecco's Modified Eagle's Medium; Corning, 10027CV). Cells were injected at rate of 0.1uL/min using a Hamilton syringe (Hamilton, Darmstadt, Germany) at pontine coordinates of 1mm posterior and 1mm lateral (right) from lambda, and 5.5mm deep. Tumors were confirmed 10 days post injection (dpi), using the Bruker Biospec 9.4 Tesla Small Animal MR Imager (Bruker Medical, Boston, MA, USA) via the Oncology Precision Therapeutics and Imaging Core at our institution.

#### FLASH Radiation

The FLASH system is based on a repurposed Varian Clinac 2100C, where the control system was modified to allow control of beam delivery on a pulse-by-pulse basis, as well as tuning of the pulse forming network to optimize Dose Per Pulse (DPP).[2] Pulse duration is < 1 μsec and pulse repetition rate is 180Hz. Prior to irradiation, mice were anesthetized using isoflurane, immobilized in a mouse holding fixture (Precision X ray Irradiation Inc, Madison, CT) and placed on a 3D printed jig, incorporating 6.4mm thick movable lead shielding which was visually aligned to irradiate the mouse hindbrain (typically 15 mm gap; Supplemental Figure 1).

For CONV, mice were individually placed at a source to skin distance (SSD) of 171cm and irradiated alongside a NIST traceable Advanced Markus Ionization Chamber (AMIC). The Clinac was set to deliver 2-4 pulses per second (1 pulse on/50-100 pulses off), adjusted periodically throughout the irradiation to compensate for drift in the DPP (average DPP=0.008 Gy), and maintain a dose rate of approximately 2 Gy/min. Irradiation was stopped manually when the AMIC reached 15Gy. A typical mouse received 1900 pulses over 8 minutes. For FLASH, mice were placed at an SSD of roughly 50 cm (Beam size 30 mm FWHM; the jig was held by the Y Jaw) and irradiated with 25-30 pulses, at a repetition rate of 180Hz (130-160 msec total duration). At this SSD, dose per pulse is approximately 0.5 Gy, the dose is uniform to within 10% within the thickness of the mouse. Beam was delivered by waiting 20 s with the klystron and electron gun on, but out of synchronization (allowing the pulse forming network to stabilize without beam delivered to the mouse) and then delivering the required number of pulses. Exact positioning of the mice and the required pulse count for each irradiation was determined on the day of irradiation by irradiating the AMIC and film at the same location. The built-in Clinac ionization chamber (and, for some experiments, EBT3 gafchromic film) was used to verify that there was no drift in beam intensity between the calibration and mouse irradiations.

**Tissue dissection and processing**

Brainstems were dissected at both day 4 (D4, scRNAseq and flow cytometry) and day 10 (D10, flow cytometry only) post-RT. Upon dissection of the brainstem, the tissue was placed in isolation media consisting of RPMI 1640 with GlutaMAX Supplement and HEPES (Gibco, 72400047) with

0.01% 50X B27 supplement (Gibco, 17504044). The tissue was then homogenized in a 15ml glass tissue grinder, filtered through a 70um filter. The pellet was resuspended in 1ml of isolation media (per brainstem) and the mixture was incubated with 100ul of myelin removal beads (Miltenyi Biotec, 130-096-733). The solution was then passed through LS+ positive separation columns (Miltenyi Biotec, 130-042-401) to deplete the myelin.

**10x genomics chromium single-cell 3' library construction**

Four samples were loaded onto a single channel of a 10x Chromium chip, one sample from each treatment group. This pooling of different samples was possible as we used Totalseq-B feature barcoding technology to multiplex our samples for computational re-identification of cells post-sequencing.[3] Viability was assessed by trypan blue exclusion assay and the number of cells taken from each sample for each channel were pooled equally so that 10,000 cells total were submitted for sequencing. This process was repeated four times for a total of 16 samples run across 4 channels. The samples were run using the Chromium Next GEM Single Cell 3' Reagent Kits v3.1 (Dual Index) with Feature Barcoding technology for Cell Surface Protein (10x Genomics). The protocol was followed according to the manufacturer's instructions.

**Single-cell RNA sequencing data pre-processing and clustering**

Barcoded reads were demultiplexed and aligned to the mm10-2020-A genome using CellRanger v5.0.1 with default parameters. The datasets were first analyzed individually and then ultimately combined and further analyzed using the Seurat R package v4.0 (Supplemental Figure 8).[4] Cells with <500 or >40,000 unique molecular identifiers (UMIs) and <400 genes were filtered out along with genes detected in <3 cells. Seurat's NormalizeData() function was used to log-normalize each RNA assay and centered log-ratio transformation (CLR) was used for the hashtag assay. After hashtag demultiplexing using Seurat's HTODemux() with default parameters, only the singlets were retained. Seurat's ScaleData() was run to regress out the number of UMIs as a source of variation after FindVariableFeatures(). After pre-processing, 33,308 cells were used for clustering. Batch correction was performed using Harmony v.0.1.1 and the corrected Harmony embeddings were used for Seurat's UMAP and Nearest Neighbor analyses.[5] Clusters were identified using the FindClusters() function and the differentially expressed genes for each cluster were determined using the FindAllMarkers() function, both in Seurat. Clusters were annotated as known immune cell types using canonical marker genes and cross-referenced using the SingleR package.[6]

**Differential gene, pathway and geneset enrichment analyses**

In evaluating the differential gene expression of FLASH compared to CONV, we first conducted two individual analyses where each RT modality was compared separately to unirradiated tumor - FLASH vs SHAM (FvS) and CONV vs. SHAM (CvS) – using Seurat's FindMarkers() function (Wilcoxon Rank Sum test). This way, the response provoked from tumor presence itself is controlled for and we only focus on the response provoked uniquely by each RT group. We then compared the common (similarly upregulated or downregulated) and different (differentially upregulated or downregulated) genes resulting from those two expression analyses in order to identify differences in FLASH vs CONV responses. Of note, when reporting the common and different genes between FvS and CvS throughout the clusters, we include only genes with an adjusted p-value <0.05 and an average log2 fold change >=|0.50|. If there were many significant genes, only the top ones are visualized with the rest included in supplementary data 4. In addition, we included an additional, more direct comparison of FLASH vs CONV for the MAC cluster volcano plot that is also included along with the two individual analyses in supplementary data 4.

For the microglia clusters, the top 50 most upregulated genes (Supplementary Data 2) were submitted to REACTOME in a treatment group agnostic manner for pathway analysis, and the top 10 most significant pathways were reported.[7] For pathway analysis done on the common upregulated genes between FvS and CvS, only genes with an adjusted p-value of <0.05 and an average log2 fold change of >=0.5 were submitted (Supplementary Data 4). Single sample geneset enrichment analysis (ssGSEA) for type 1 interferon response (IFN1) was performed using the Escape R package on the cells from the cluster under analysis with the REACTOME_INTERFERON_ALPHA_BETA_SIGNALING geneset.[7,8] Treatment groups were evaluated separately and compared two at a time using a non-parametric Mann Whitney U test. Significance was defined as p value ≤0.05. Flow cytometry statistical methods are reported in the main manuscript.

**Flow Cytometry**

After the tissue was processed, the pellet was first incubated with a 1:100 blocking solution containing TruStain FcX (anti-mouse CD16/32; Biolegend, 156603) and isolation media before antibody staining. Cells were stained with an antibody cocktail containing isolation media and anti-mouse 7AAD viability solution (420403), anti-mouse CD45 (157617), anti-mouse CD11B (101263), anti-mouse CD163 (155305), anti-mouse CD206 (Mrc1, 141727), anti-mouse CD11C (117327), anti-mouse I-A/I-E (MHC-II, 107619), anti-mouse CD3 (100229), anti-mouse CD8 (126617), anti-mouse CD4 (eBioscience, 35-0042-80), anti-mouse GR1 (Ly6G/Ly6C, 108409), anti-mouse TMEM119 (eBioscience, 12-6119-82), anti-mouse CD19 (115540), and anti-mouse IFNAR1 (127325). All antibodies were purchased from BioLegend unless otherwise noted and used at a 1:100 dilution. Fluorescence-minus-one (FMO) samples were generated at the same time and used to determine appropriate gating cutoffs for each flow cytometry run. All samples were run using the Cytek Aurora at our institution's Center for Translational Immunology and analyzed using the FlowJo software (v.10.8.2). Methods for statistical analysis of the flow cytometry data are reported in the main manuscript.
